# KDM6B interacts with nucleo-adhesome components CSRP2 and TGFB1I1 to regulate EMT

**DOI:** 10.64898/2026.08.24.737021

**Authors:** Durand Jules, Montagnon Frederic, Jaramillo Ortiz Sarahi, Schaeffer-Reiss Christine, Herfs Michael, Nokin Marie-Julie, Diane Bruyère, Fabienne Perin, Pallandre Jean René, Borg Christophe, Peigney Anne, Overs Alexis, Lupien Mathieu, Guittaut Michael, Hervouet Eric, Delage-Mourroux Regis, Peixoto Paul

## Abstract

The methyltransferase EZH2 (Enhancer of Zest Homolog 2) and the demethylase KDM6B (Lysine Demethylase 6B) have been associated with epithelial to mesenchymal transition (EMT) and poor prognosis in various cancers(1–9). These enzymes methylate and demethylate H3K27me3 and regulate distinct sets of genes controlling EMT induction, despite having opposite catalytic activities(10). This could be due to their recruitment or the modulation of their activity by partner proteins on specific *loci*. This work sought to identify proteins associated with chromatin and interacting with EZH2 or with KDM6B during EMT. To do so, co-immunoprecipitation and mass spectroscopy was used under TGFβ (Tumor growth factor β) and TNFα (Tumor necrosis factor α) treatment to induce EMT in A549 lung cancer cells. Surprisingly, numerous proteins related to focal adhesions were identified to interact with EZH2 or KDM6B. These proteins are part of a nuclear protein interaction network previously described as nucleo-adhesome. Among these proteins, TGFB1I1 (transforming growth factor induced peptide 1) and CSRP2 (cysteine and glycine rich protein 2) were further confirmed to interact with KDM6B in the nucleus and even more so during EMT. The target genes of these complexes were then sought by knocking down KDM6B, TGFB1I1 or CSRP2. Three genes (coding Integrin alpha 5, Laminin y2 and Matrix Metalloproteinase 9) were confirmed to be regulated by KDM6B, TGFB1I1 and CSRP2. These findings may have clinical relevance, as immunohistochemistry analyses performed on a cohort of lung cancer patients revealed increased nuclear localization of TGFB1I1 and CSRP2 in cells undergoing EMT.

## INTRODUCTION

EMT governs metastases formation which remains strongly associated with poor prognosis in many different cancers. In lung cancers for example, 5 years overall survival (OS) drops from 47% in stage 1 cancers (localized tumors) to 5% in stage 4 cancers (metastatic cancers). Despite large evidence that EMT is an epigenetic-driven phenomenon(11–13), the precise mechanisms governing gene expression reprogramming in EMT are still largely unresolved. We and other teams previously identified H3K27 methylation as a key histone mark in EMT regulation(14,15) and showed an important role of the enzymes EZH2 and KDM6B, which respectively methylate and demethylate H3K27. These enzymes contribute in the specific epigenetic control of EMT in various cancers(16,17) including non-small cell lung carcinoma (NSCLC)(10,18). Expression of EZH2 and of KDM6B has been associated to aggressiveness or metastasis in various cancers (1–9). EZH2 is part of the repressive PRC2 complex (polycomb repressor complex 2) which includes various components proteins. Embryonic Ectoderm Development (EED) and JARID2 (Jumonji, AT rich interactive domain 2) are two well-described components of PRC2 with a shown function in recruiting EZH2 towards EMT related genes during TGFβ EMT induction(19,20). For example, JARID2 has been shown to recruit the PRC2 complex on the promoter of the E-Cadherin encoding gene *CDH1*, leading to its repression by H3K27 methylation during TGFβ induced EMT in the NSCLC cell line A549 and the colon cancer cell line HT29(20). Similarly, KDM6B is also recruited on different genes during EMT(10,21) by mostly unknown processes, and its partner proteins are poorly described in comparison of those of EZH2. To further understand the fine tuning of H3K27 methylation during EMT we investigated the proteins interacting with EZH2 and KDM6B during TGFβ1/TNFα (transforming growth factor beta 1 and tumor necrosis factor alpha) induction of EMT in A549 cells (lung adenocarcinoma) which could explain their recruitment on specific loci to regulate EMT-related genes. Using a co-immunoprecipitation/mass spectrometry (co-IP/MS) approach adapted from the Rapid Immunopreciptation and Mass spectrometry of Endogenous proteins (RIME) protocol(22), we identified multiple proteins related to integrin cell adhesion (adhesome components) that co-precipitated with EZH2 and KDM6B. Considering it was recently described that adhesome proteins can locate to the nucleus and regulated gene expression as part of a large network termed nucleo-adhesome(23), we then focused our analysis on two of these proteins: TGFB1I1 (Transforming growth factor beta 1 induced transcript 1) and CSRP2 (Cysteine and Glycine Rich Protein 2). We found that these proteins interact with KDM6B in the nucleus to regulate epigenetic targets of KDM6B that promote a mesenchymal phenotype. These results deliver proof of concept that nucleo-adhesome components may have a role in regulating adhesion and EMT by exerting control over epigenetic regulations.

## MATERIAL AND METHODS

### Cell culture, treatments and transfection

The A549 (NSCLC) cell line was obtained from Dr. Christophe Borg (INSERM UMR1098, Besançon, France). Stable A549 cell lines overexpressing EZH2 or HA-KDM6B were obtained as previously described(10). Cells were grown in DMEM 1 g/L glucose (L0066, Dominique Dutscher, Brumath, France) supplemented with 10% fetal bovine serum 10% (S1810, Dominique Dutscher), penicillin-streptomycin (50 U/mL) (L0018, Dominique Dutscher) and amphotericin B (1.25 µg/mL) (P11-001, PAA). Cells were cultured at 37°C in 5 % CO_2_ and routinely used at 70−80% confluence. When indicated, cells were treated for 48 h using 4 ng/mL TGFβ (100-21, Peprotech) and 20 ng/mL TNFα (300-01A, Peprotech, Rocky Hill, NJ, USA). siRNA CSRP2 (Dharmacon, Lafayette, CO, USA, L-011244-00-0005), siRNA KDM6B (Dharmacon, L-023013-01-0005) and siRNA TGFB1I1 (Dharmacon, L-006565-00-0005), were transfected using lipofectamine 2000 (Invitrogen, Carlsbad, CA, USA, 11668019) according to the manufacturer’s protocol.

### Co-IP/MS and ChIP

A549, A549-EZH2 or A549-HA-KDM6B were cultured with or without TNFα/ TGFβ treatment and recovered by trypsinization and cells were centrifuged for 3 min at 500 g, washed with HBSS (Hank’s Balanced Salt Solution) and centrifuged again for 3 min at 500 g. For co-IP/MS analysis Crosslinking was performed by resuspending the cells in PBS (Phosphate Buffered Saline) with 0.32µg/µL Disuccinimidyl glutamate (0.32 µg/µL, Sigma-Aldrich-Aldrich, St. Louis, MO, USA, 80424) and incubating for 22 min under agitation at room temperature followed by the addition of 1% formaldehyde (Sigma-Aldrich-Aldrich, F8775) and an additional 8 min under the same conditions. For ChIP analysis only the fixation with formaldehyde was performed. Quenching was performed by adding glycine (0.125 M final, Euromedex, Strasbourg, France, 26-128-6405-C) for 5 min keeping cells under agitation. Cells were then centrifuged for 4 min at 500 g and 4°C, washed twice with cold PBS containing a protease inhibitor cocktail (PIC, 1 X, Bimake, B14001) before being resuspended and incubated for 10 min with agitation at 4°C in cell lysis buffer (Tris/HCl pH 8 20 mM, KCl, 85 mM, NP40 0.5%, PIC 1 X). To recover the nuclei, the cells were then subjected to mechanical lysis in a homogenizer (DWK Life sciences, Wertheim, Germany, Wheaton dounce, “Tight”). After 5 min centrifugation at 1700 g and 4°C, the supernatant was removed and the nuclei were washed twice with buffer (Tris/HCl pH 7.6 10 mM, EDTA 1 mM, SDS 0.1%, PIC 1 X) without resuspension. Nuclei were then resuspended in the same buffer and chromatin was fragmented by ultrasound (Covaris, Woburn, MA, USA, E220 Sonicator program: 15 min, 7°C, 115 W, duty cycle 2%, cycle per burst 200). Finally, chromatin concentration was assayed using the Qubit 3 fluorometer (Invitrogen, Q33216) using a Qubit dsDNA BR Assay kit (Invitrogen, 32850).

Immunoprecipitations were performed using an IP-Star® Compact Automated System (Diagenode, Liège, Belgium, BO3000002). Co-IP/MS was performed with 25 µg of chromatin and ChIP with 5µg. Chromatin was immunoprecipitated by a 6 h incubation with the appropriate antibody (**Table 1**). Twenty microliters of G protein-coupled magnetic bead suspension (Active Motif, Carlsbad, CA, USA, 53033) previously washed in 20 mM Tris-HCl pH 8 buffer, 167 mM NaCl, 0.01% SDS, 1.1% Triton X100, 1.1 mM EDTA, 0.1 X PIC, 0.1% BSA were added and incubated for 2h. Finally, beads bound to complexes of interest were washed 3 times with Tris-HCl buffer pH 8 20 mM, NaCl 167 mM, SDS 0.01%, Triton X100 1.1%, EDTA 1.1 mM, PIC 0.1 X. Samples were stored at 4°C during all these steps.

### Proteomic analysis

#### Sample preparation

Protein concentration was estimated using a Pierce 660-nm Protein Assay Kit (Thermo Fisher Scientific Waltham, MA, USA). Samples were diluted with a 0.1 M ammonium bicarbonate (ABC) solution to achieve a final volume 50 µL, containing 20 µg of proteins. Reduction was performed using Dithiothreitol (DTT) to reach a final concentration of 12 mM, incubating at 37°C for 30 min. Protein alkylation was performed for 30 minutes at room temperature in the dark using 175 mM iodoacetamide (IAM) in a 0.1 M BCA solution to obtain a final IAM concentration of 40 mM. SP3 digestion was performed using a 1:1 mixture of beads A (Sera-Mag Speed, Thermo Fisher Scientific, 45152105050250) and beads B (Sera-Mag Speed, Thermo Fisher Scientific, 65152105050250). After performing three washing steps with H_2_O, the combined beads were added to the samples at a beads-to-protein ratio of 10:1 for each type of bead, resulting in an overall ratio of 20:1 for the combined beads. For effective protein binding to the beads, acetonitrile (ACN) was added to have a final concentration of 50%, and the mixture was allowed to incubate for 18 minutes. The beads were washed twice with 80% ethanol (EtOH) and once with 100% ACN before resuspension in 95 μL of ABC. Trypsin/Lys-C (Mass Spec Grade mix, Promega, Madison, WI, USA) was added at an enzyme-to-protein ratio of 1:50 and the proteins were digested overnight at 37°C, 1,000 rpm. Digestion was stopped by the addition of trifluoroacetic acid (TFA) (final pH < 2). After peptide recovery, the samples were dried using a vacuum centrifuge and resuspended in 2% ACN and 0.1% formic acid (FA).

### LC–MS/MS analysis

Samples were analyzed using a nanoflow liquid chromatography system (nanoElute, Bruker Daltonics) coupled online to a trapped ion mobility spectrometry-quadrupole time-of-flight mass spectrometer (timsTOF Pro 2, Bruker Daltonics Billerica, MA) equipped with a CaptiveSpray source operating in positive ion mode. Peptides (4 µL) were loaded onto an Acclaim™ PepMap™ 100 C18 precolumn (100 µm × 20 mm, 5 µm particle size) and subsequently separated on a reversed-phase Aurora C18 column (20 mm × 180 µm, 1.6 µm particle size; IonOpticks). The solvent system consisted of 0.1% AF in water (solvent A) and 0.1% FA in acetonitrile (solvent B). Chromatographic separation was performed using a gradient starting at 2% solvent B, ramping to 15% over 19 minutes, followed by increase to 23% over 7 minutes, 30% over 4 minutes, and finally reaching 85% in 3 minutes, where it was held for an additional 7 minutes. The flow rate was maintained at 300 nL/min, and the column temperature was kept constant at 50°C. Samples were injected in randomized order, with a blank injection between each sample.

Mass spectrometry data were acquired using the PASEF method, with 10 PASEF MS/MS scans over an *m/z* range of 100–1700 and a total cycle time of 1.88 s. The ion mobility coefficient (1/K₀) was set between 0.7 and 1.25 Vs cm⁻², with both ramp and accumulation times set at 166 ms. Singly charged precursors were excluded based on their position in the m/z–ion mobility plane, and precursors that reached a target value of 17,000 a.u. were dynamically excluded for 0.4 min. The quadrupole isolation width was set between 2 and 3 m/z, depending on the precursor m/z. Collision energies ranged from 20 to 52 eV. The source temperature was maintained at 180°C, with a spray voltage of 1.6 kV. TIMS, MS operation, and PASEF were controlled and synchronized using OtofControl 6.0 software (Bruker Daltonik).

### Mass spectrometry data processing

Raw MS files were converted into Mascot Generic Format (MGF) files and subjected to a Mascot (version 2.6.2, Matrix Science) search using a target-decoy strategy to assess the false discovery rate (FDR). The human database for the search was generated in-house and included human entries from UniProtKB/Swiss-Prot, totaling 20,419 entries as of September 12, 2019. The search parameters were as follows: trypsin was set as the cleavage enzyme, allowing a maximum of one miscleavage. Methionine oxidation and acetylation of protein N-termini were set as variable modifications, while cysteine carbamidomethylation was established as a fixed modification. The peptide mass tolerance was set to 15 ppm, and the MS/MS mass tolerance was set to 0.05 Da.

Proline(24) was used for validation and protein identifications were set as follows: a minimum length of seven amino acids, a score of ≥20, a pretty rank of ≤1, and a FDR below 1% at both the PSM and protein levels. For label-free quantification, peptide abundances were extracted using an m/z tolerance of 15 ppm. Alignment of the LC-MS runs was performed using Loess smoothing. Cross-assignment of peptide ion abundances was conducted among the samples and controls, using an m/z tolerance of 5 ppm and a retention time tolerance of 60 s. Protein abundances were computed using the median ratio fitting of unique peptide abundances, normalized at the peptide level using the median.

Differential data analysis was conducted using the open-source ProStaR software (version 1.34.6)(25). Only proteins identified five times in at least one condition (i.e., in all replicates of a condition) were retained for the quantitative study. The intensities were log_2_-transformed and associated with their respective conditions. Protein abundances were normalized using 0.15% quantile centering. Imputation of missing values was performed by approximating the lower limit of quantification using the 2.5% lower quantile of each replicate intensity distribution (referred to as ‘det quantile’). A Limma moderated t-test was applied to the dataset for differential analysis. The calibration of p-values was corrected using the adapted Benjamini-Hochberg method to adjust for multiple testing, and differentially expressed proteins were identified using a p-value threshold. Given the small number of differentially expressed proteins observed, the FDRs could not be set at the conventional 1%, and were instead set at the lowest relevant value: IP KDM6B vs IgG 2,19 % FDR (max p value 0,000631) ; IP KDM6B untreated vs TGFβ/TNFα 6,06 % FDR (max p value 0,000794) ; IP EZH2 vs IgG 4,24 % FDR (max p value 0,000794) ; IP EZH2 untreated vs TGFβ/TNFα 2,08 % FDR (max p value 0,000631).

### Subcellular fractionation

Cells were washed with ice-cold PBS, then scraped Buffer A (10mM HEPES pH 7.9, 10mM KCl, 1.5mM MgCl_2_, 0.34M Sucrose, 10% Glycerol, 0.1% Triton X-100, PIC 1X). The suspension was incubated for 5 minutes at 4°C under gentle rotation then centrifuged at 1300g for 4 minutes at 4°C. The first supernatant was used as cytosolic fraction, and the nuclear pellet was washed twice Buffer A and centrifuged under the same conditions. The pellet was then resuspended in RIPA buffer (Tris HCl 50 mM, NaCl 150 mM, 0,1% SDS, 1% Triton X100, 0,5% DOCA, pH 7,4, PIC 1X), sonicated following the same condition as for the co-IP/MS experiment and used as nuclear fraction.

### Western-blot and dot blot

Protein concentration in nuclear or cytoplasmic fractions was assessed using the DC protein Assay (BioRad, Hercules, CA, USA, 5000112). Protein samples were separated on 7.5% polyacrylamide gel (TGX Stain Free, 161-0181, BioRad) in migration buffer (Tris 25 mM - Glycine 196 mM pH 8.3, SDS 0,1%). Migration was performed at 100 V in the concentration gel, then at 150 V in the separation gel. Proteins were then transferred to a PVDF membrane (1620177, BioRad) previously activated in a 100% ethanol bath for 10 min. Transfer was carried out using the Trans-Blot Turbo system (1704150, BioRad) according to the predefined “Mixed MW” program. Total proteins were revealed on the membrane using Stain-Free technology (BioRad) and ChemiDoc XRS+ (BioRad, California).

Dot blots were performed by eluting IP products in 20µL Laemmli 2X (Tris 125 mM pH 6.8, SDS 4%, Glycerol 20%, β-mercaptoéthanol 10%) and heating at 95°C 10 min. For each dot, 2µL of elution was loaded on a nitrocellulose membrane then dried overnight at RT.

For both western and dot blot, the membrane was saturated in a solution of TBS-T (Tris 20 mM NaCl 150 mM, pH 7.5, Tween 20 0.1%) containing 5% BSA for 1 h with agitation, then incubated overnight at 4°C with agitation in a solution of TBS-T 5% BSA containing the primary antibody of interest (**Table 1**). The membrane was then washed 3 times for 10 min in TBS-T with agitation, then incubated for 1 h with agitation at room temperature in TBS-T containing a secondary anti-mouse 1/10,000 (BI-2413C) or anti-rabbit 1/10,000, (Abliance, Compiègne, France, BI-2407) antibodies coupled to Horseradish Peroxidase (HRP). After 3 x 10 min washes in TBS-T with agitation, HRP activity was revealed after incubating the membrane for 5 min in the dark with 500 µL Clarity Western ECL Substrate reagent (1705061, BioRad). The luminescence emitted by the HRP was then quantified using ChemiDoc XRS+ (BioRad) and Image Lab software.

### Immunofluorescence, Proximity ligation assay and confocal microscopy

Cells were cultured and treated on cover slides then washed with cold PBS, fixed with 4% PFA in PBS for 10 min on ice and permeabilized using PBS 0.1% triton-X100 10 min on ice. The saturation was performed using Duolink Blocking Solution (Merck, Darmstadt, Germany, DUO82007) for 1 h at 37 °C. Cover slides were then removed and incubated overnight at 4 °C with primary antibodies diluted in 1% BSA (**Table 1**) before being washed three times with PBS-Tween 0.1% (PBS-T) under agitation. For the IF, a secondary antibody Alexa Fluor 488 goat anti-mouse or rabbit (Thermo Fisher Scientific, A11001 and A11008) was incubated for 1 h at 37 °C. After 3 washes with PBS-T the nuclei were stained in 10µM DAPI (Euromedex, 1050-A) in PBS for 10 min under agitation, before the cells were mounted using fluormount medium (F4680, Merck). For the PLA, the Duolink (DUO92101, Merck) was used in accordance with the manufacturer’s instructions. Image acquisition using a Zeiss LSM800 AiryScan laser scanning confocal microscope with a 63X objective. Dots were counted using a homemade CellProfiler(26) pipeline and R script.

### RNA extraction and reverse transcription

RNA extraction was performed using NucleoSpin RNA Plus (Macherey-Nagel, Düren, Germany, 740984.250) according to manufacturer’s instructions and reverse transcription was performed with PrimeScript RT Reagent Kit (Takara, Kusatsu, Shiga, Japan, RR047B) according to manufacturer’s instructions.

### RNA sequencing and analysis

RNA sequencing was performed by the MGX platform (Montpellier GenomiX, University of Montpellier), the libraries were constructed using Illumina’s Stranded mRNA Prep Ligation kit (Illumina, San Diego, CA, USA, 20040534) and sequenced on an SP flow cell single-read 100 nucleotides with Illumina’s NovaSeq 6000 System.

Reads were trimmed with the trimmomatic software(27), pseudo aligned with kallisto(28) then abundances were normalized with DESeq2(29) all using default parameters. Downstream analyses were performed using R.

### ATAC-seq

ATAC libraries were created using Active motif ATAC-Seq Kit (Active motif, 53150) and sequenced using Illumina’s Novaseq6000 and SP flowcell paired-end 50 nucleotides. All samples were processed using the ENCODE ATAC-seq pipeline to call peaks and IDR. Normalization, Fold changes and p-value calculations were performed using Diffbind with default parameters.

### qPCR

For ChIP as well as cDNA obtained from reverse transcription, qPCR reactions were performed using the Green® Premix Ex-Taq™ kit (Takara, RR420L) using the sense and antisense primers listed in **Table 2**.

### Wound healing assay

Following siRNA transfection and TGFβ/TNFα as described previously, cells were collected by trypsinization and seeded at a confluence of 150 000 cells per well and left to adhere overnight. Scratches were performed with pipette tips and followed by two washes with 37°C HBSS, then new medium with or without TGFβ/TNFα was added. Four individual experiments were conducted, which included two pictures per well and two wells per condition. Migration was monitored for 8 h and wound surface area was measured using ImageJ.

### Tissue specimens, immunohistochemistry and immunostaining assessment

Paraffin-embedded tissue samples from 47 patients with non-small cell lung cancer were retrieved from pathology archives [in collaboration with the Tissue Biobank of the University of Liège (Liège, Belgium)]. The original diagnosis was confirmed by experienced pathologists. Based on Vimentin expression in tumor cells, samples were categorized into EMT-positive and EMT-negative groups according to previous work(10,15,30). Ethical approval for the study was granted by the Ethics Committee of the University Hospital of Liège (#2024-262).

Technically, after tissue sectioning, deparaffinization, and rehydration, antigen retrieval was performed using either 10mM citrate buffer (Agilent, Santa Clara, CA, USA, for TGFB1I1 staining) or target retrieval buffer (Agilent, for CSRP2 staining) for 10 min at 126°C. Sections were then incubated with 3% hydrogen peroxide for 20 min at room temperature (RT) to inhibit endogenous peroxidase activity. To prevent non-specific antibody binding, tissue sections were treated with an animal-free blocking solution (Cell Signaling Technology, Danvers, MA, USA) for 10 min at RT, before being exposed to the primary antibody [monoclonal anti-Vimentin (clone V9, Ventana Medical Systems, ready to use), polyclonal anti-TGFB1I1 (10565-1-AP, Thermo Fisher Scientific, 1:100) or anti-CSRP2 (clone E8R5N, 28601, Cell Signaling Technology, 1:200)] for 1 h at RT. After a washing step, an anti-mouse or anti-rabbit HRP secondary antibody (EnVision+ Single Reagent, Agilent) was added for 30 min at RT. Positive cells were visualized using 3,3’ diaminobenzidine tetrachloride (DAB). Slides were finally counterstained with hematoxylin, dehydrated and mounted. Mouse or rabbit control IgG (Santa Cruz Biotechnology, Dallas, TX, USA) were used as negative control.

Anti-TGFB1I1 and anti-CSRP2 immunolabelled tissues were evaluated independently by senior investigators using a semi-quantitative scoring system. Staining intensity was scored as follows: 0 = negative, 1 = low, 2 = moderate, and 3 = strong. The extent of staining was scored based on the percentage of positive cells: 0 = <5%, 1 = 6–33%, 2 = 34–66%, and 3 = >67%. For each case, the entire malignant lesion was assessed. As previously described(31,32), the scores for intensity and extent were multiplied to obtain a final score. Notably, the presence or absence of cellular patches (defined as clusters containing at least 50 positively stained cells) showing nuclear immunoreactivity was also recorded.

## RESULTS

### EZH2 and KDM6B are associated with specific protein partners in EMT

Since both EZH2 and KDM6B, with opposite catalytic activity, are frequently overexpressed in EMT and recruited on different genes, we hypothesized that these proteins could be controlled by specific protein partners during this process. To explore this hypothesis, we performed co-IP/MS experiments in A549 cells stably expressing EZH2 or HA-KDM6B treated or not with TGFβ1 and TNFα to induce EMT, as the crosstalk between these two cytokines in the tumoral microenvironment(33) has previously been reported to enhance EMT in the A549 model(34–36). HA-tagged protein strategy was selected due to a lack of efficient antibodies to immunoprecipitate endogenous KDM6B protein. The effects of EZH2 or KDM6B overexpression on EMT an efficiency of EMT induction by TGFβ/TNFα were measured validated using both microscopy and RT-qPCR (**Supp Figure 1 A-C**). The co-IP/MS protocol includes protein-DNA crosslinking as well as nuclear enrichment which facilitates the identification of DNA-interacting complexes. Correct shearing of the chromatin as well as specificity and efficiency of IP against EZH2 and HA-KDM6B were first validated as illustrated in **Supp Figure 1 D and E.** After MS analysis, only proteins identified in 5 out of 5 independent experiments were considered as putative partners.

**Fig. 1.**
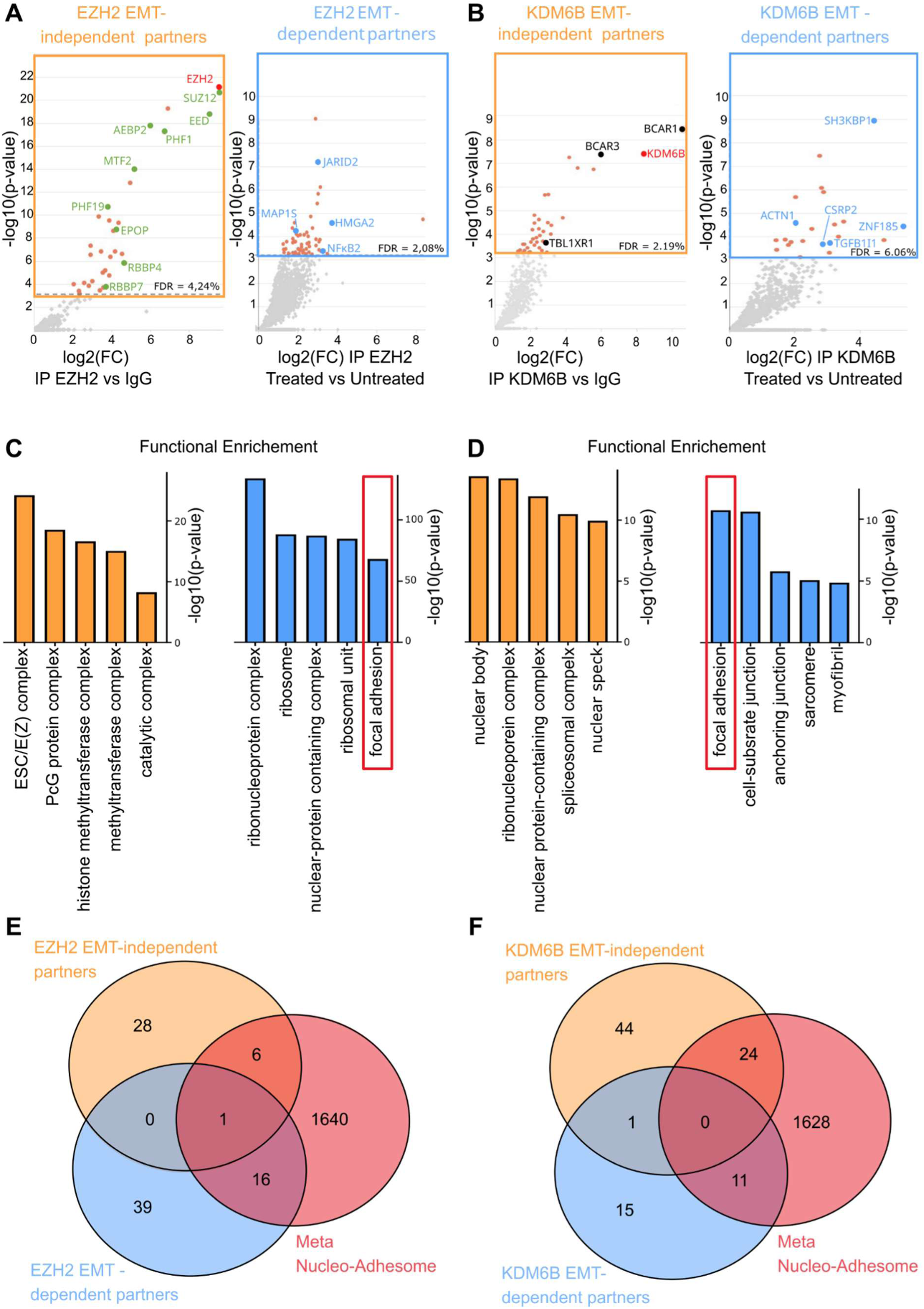
Co-IP-MS identification of the EZH2 and KDM6B partners in A549 induced or not in EMT. A549-EZH2 or A549-HA-KDM6B cells were treated or not with TGFβ/TNFα and EZH2 and KDM6B protein partners were investigated by Co-IP-MS analysis. (A) Volcano plots for EZH2 partners in basal condition (IP EZH2 vs IgG) and during EMT (IP EZH2 treated vs untreated). (B) Volcano plots for KDM6B partners in basal condition (IP EZH2 vs IgG) and during EMT (IP EZH2 treated vs untreated). (C) Gene ontology analysis of TGFβ/TNFα independent and dependent partners of EZH2. (D) Gene ontology analysis of TGFβ/TNFα independent and dependent partners of KDM6B. (E) Venn diagram overlapping EZH2 partners with the nucleo-adhesome set of protein. (F) Venn diagram overlapping KDM6B partners with the meta-nucleo-adhesome set of protein.

As expected, EZH2 and KDM6B were among the most significantly enriched proteins in the IP versus IgG comparisons, confirming the specificity and efficiency of the immunoprecipitation. Almost no partners were found significantly less co-immunoprecipitated with EZH2 or KDM6B after EMT induction, (negative fold changes in IP EZH2/KDM6B treated versus untreated conditions) suggesting the complexes formed during EMT were either newly formed or formed by the addition of new components to pre-existing complexes. This observation led us to distinguish, for EZH2 and for KDM6B, “EMT-independent partners” (significantly enriched in IP EZH2/KDM6B vs IgG, untreated condition) from “EMT-dependent partners” (significantly enriched in IP EZH2/KDM6B treated versus untreated conditions).

Regarding EZH2, 35 EMT-independent partners were identified and included SUZ12 (Suppressor of zeste 12 protein homolog), EED, AEBP2 (Adipocyte Enhancer-Binding Protein 2), RBB4 (Retinoblastoma-binding protein 4), RBB7 (Retinoblastoma-binding protein 7), all well-established components of the PRC2 complex. Moreover, 56 EMT-dependent partners were found for EZH2 (**Figure 1A**) including JARID2, a non-essential PRC2 which was already shown to recruit EZH2 towards epithelial genes during TGFβ EMT induction of A549 cells(20). Similarly, 69 proteins were specifically interacting with HA-KDM6B in basal condition and 26 additional EMT-dependent partners (**Figure 1B**) were identified. Only 4 proteins were common between EZH2 IP and KDM6B IP sets (SH3 domain-containing kinase-binding protein 1 SH3K1, Sphingosine-1-phosphate lyase 1 SGPL1, Alpha-actinin-1 ACTN1 and Paraspeckle component 1 PSPC1) suggesting that these proteins with opposite catalytic activities are, as expected, included in distinct complexes. Detailed lists of these interactomes are available on **Supp Tables 1-4**. Gene Ontology (GO) enrichment analysis of these four different sets of proteins was first performed to validate the approach using the basal sets and to identify any overrepresented characteristics among EMT-dependent partners (**Figure 1C and D**). EMT-independent partners of EZH2 showed an expected enrichment in proteins of the polycomb complex and histone methyltransferase activity, whereas EMT-independent partners of KDM6B showed enrichment in nuclear body, nuclear speckles localization and components of the spliceosome complex. These results are consistent with efficient nuclear enrichment in our co-IP/MS protocol and also suggest that KDM6B may play a role in RNA splicing, which could be of interest considering the recent studies involving H3K27 methylation in RNA splicing(37–39). Notably, EMT-dependent partners of KDM6B showed an enrichment in focal adhesion components, also found with EMT-dependent EZH2 partners in lower proportions. Proteins such as CSRP2, TGFB1I1, ACTN1 or PARVA (Parvin alpha) are adhesome constituents, that interact directly or indirectly with focal adhesions and can act as Actin bundling proteins or signal transducers. Their interaction with KDM6B is consistent with the description by Byron *et al*.(23) of a nucleo-adhesome, a large network of proteins belonging to the adhesome that can interact together and are also located in the nucleus. Byron *et al.* also showed that TGFB1I1 is a member of the nucleo-adhesome capable of regulating gene expression. Furthermore both TGFB1I1(40–43) and CSRP2(44–47) have been extensively shown to locate in the nucleus as well as focal adhesions in various context to either act as co-regulators, Actin bundling proteins or signal transducers.

Comparison of our putative EZH2 or KDM6B partners with the meta nucleo-adhseome identified by Byron et.*al.* (**Figure 1E**) reveals that 23 EZH2 partners belong to the meta nucleo-adhesome, 17 of which are EMT-dependent. Among KMD6B partners, 35 belonged to the meta nucleo adhesome, 24 of which were EMT independent (**Figure 1F**). Physical interaction analysis using the String database further hints toward adhesion related partners interacting with EZH2 and KDM6B (**Supp Figure 2**). For EZH2, this analysis showed three major clusters of interacting partners: PRC2 components, RNA metabolism related proteins and focal adhesion related proteins. For KDM6B, a large network of RNA splicing and RNA metabolism wash highlighted. Focal adhesions related proteins included three directly interacting partners: BCAR1, BCAR3 (Breast cancer anti-estrogen resistance protein 1 and 3) and NEDD9 (Enhancer of filamentation 1). Interestingly, a last cluster organized around RNF20 RNF 40 (RING finger protein 20 and 40, also known as E3 ubiquitin-protein ligase BRE1A and B), two proteins that form a complex responsible for H2BK120 ubiquitination.

**Fig. 2.**
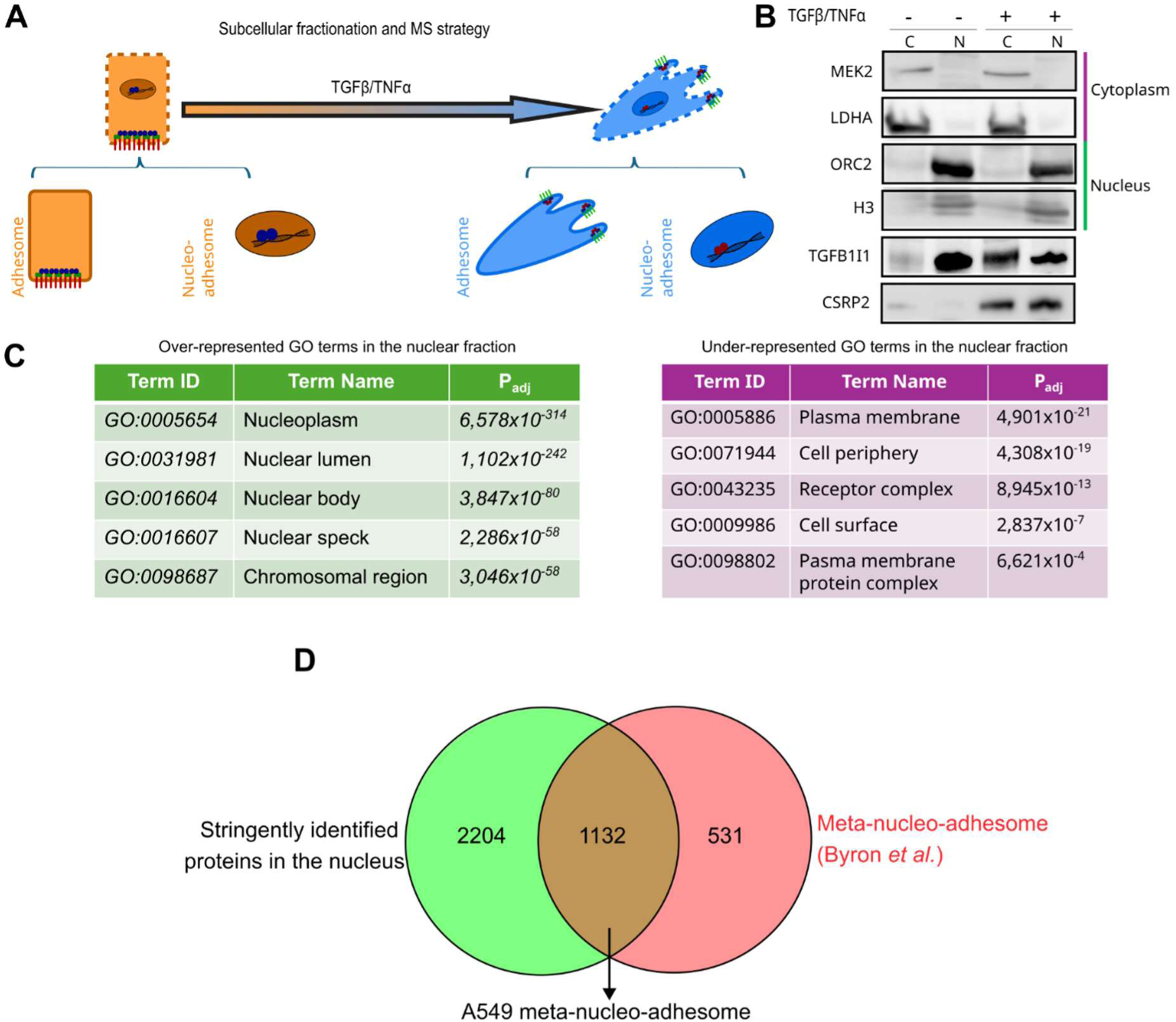
Identification of the nucleo-adhesome components of A549 cells. (A) Following subcellular fractionation, protein composition of nuclear and cytoplasmic fractions of A549 cell induced on not in EMT using TGFβ/TNFα was analyzed using LC-MS/MS. (B) Quality of the fractionation was verified using western blotting and MEK2 and LDHA, ORC2 and H3 as markers of the cytoplasm or nucleoplasm. (C) Positive and negative gene ontology analysis of the proteins identified in the nucleic fractions. (D) Stringently identified proteins in the nuclear fraction were compared to the components of the meta-nucleo-adhesome described by Byron *et al*.

### TGFB1I1 and CSRP2 illustrate a broader change in nucleo-adhesome composition during EMT

To confirm the presence of a nucleo-adhesome and assess its composition in our model, we performed subcellular fractionation followed by proteomic analysis in A549 induced or not in EMT with TGFβ/TNFα (**Figure 2A**). Correct fractionation was validated by western-blotting using cytoplasmic markers MEK2 (Dual specificity mitogen-activated protein kinase kinase 2) and LDHA (L-lactate dehydrogenase A chain), and nuclear markers ORC2 (Origin Recognition Complex subunit 2) and histone H3 (**Figure 2B**). Positive and negative GO term enrichments were also performed on the proteins identified in the nuclear fraction by LC/MS (**Figure 2C**). These analyses confirmed that the fractionation enriched the nuclear fraction in nucleoplasm and chromatin proteins while depleting cytoplasm and plasma membrane proteins. Only proteins detected in the nuclei fraction in 5 out of 5 replicates of one or both conditions (TGFβ/TNFα treated or not) were considered for the subsequent analyses. First, we compared the proteins identified in the nucleus in our analysis with the meta-nucleo-adhesome set of proteins which resulted in about two thirds of the latter being also identified in our analysis (**Figure 2 D)**. These results confirm the nuclear localization of these proteins and suggest that the nucleo-adhesome composition may be partly cell-specific.

Next, we sought to assess any changes in composition of the nucleo-adhesome during EMT. To do so, proteins detected in the nuclear fraction had their abundance compared between the two conditions using multiple t-tests then significantly Differently Abundant proteins (DAP) were selected using a 5% FDR (Benjamin-Hochberg method). Results show that 144 proteins were significantly overrepresented and 61 were underrepresented in EMT-induced nuclear fractions compared with basal nuclear fractions. These DAP were then compared with the A549 meta-nucleo-adhesome (**Figure 3A and B)**. Results show that 84 proteins, a third of the identified DAPs, belong to the meta-nucleo-adhesome, of which 59 were overrepresented and 25 were underrepresented nuclear fractions after EMT induction. Amongst the overrepresented meta-nucleo-adhesome proteins, three were EMT-dependent KDM6B partners identified by our co-IP/MS: CSRP2, ACTN1 and SH3KBP1.

**Fig. 3.**
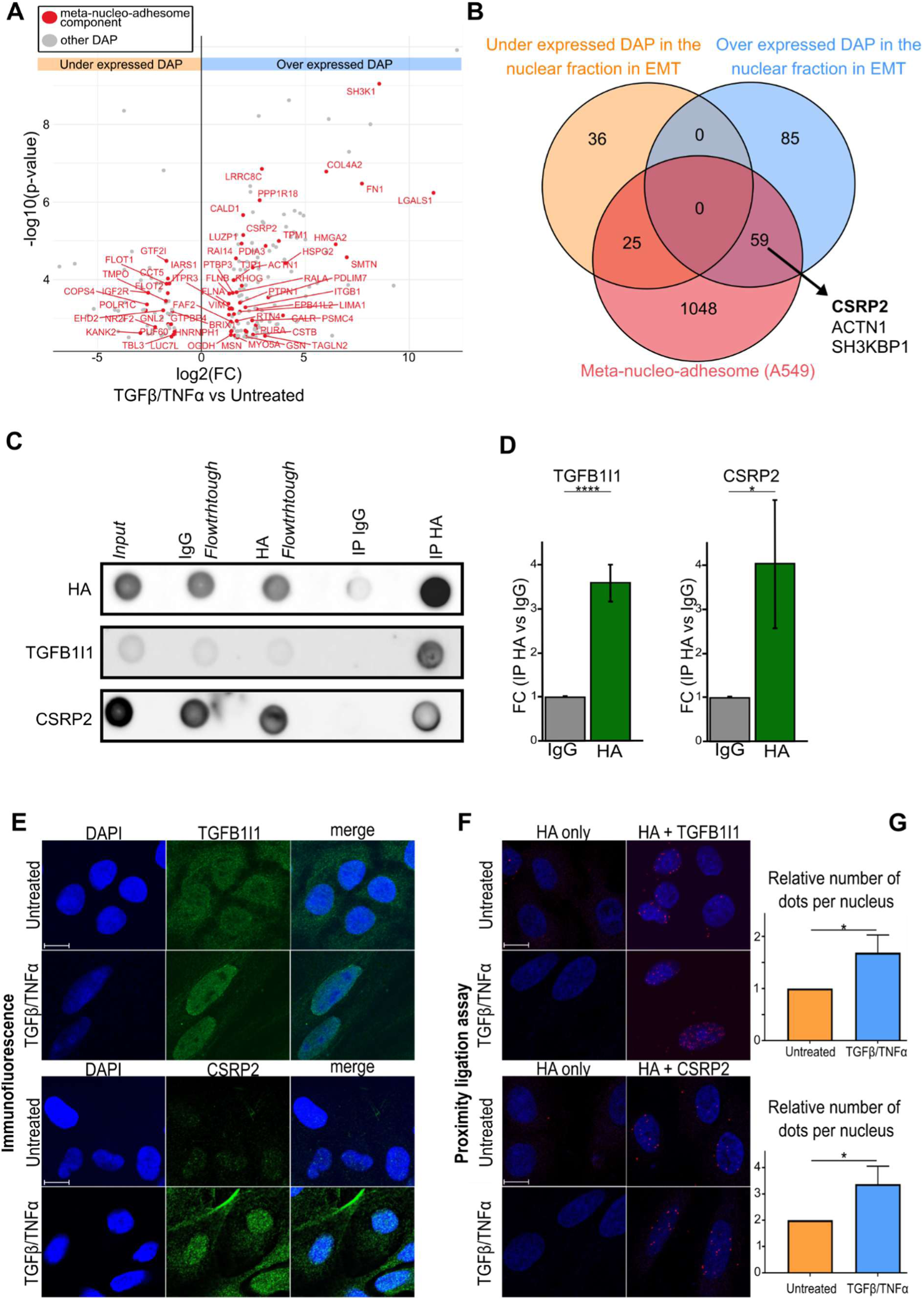
TGFB1I1 and CSRP2 illustrate a broader remodeling of the nucleo-adhesome during EMT. Following sub-cellular fractionation and MS, the abundance of each protein identified in the nuclear fraction was compared between untreated and TGFβ/TNFα treated conditions. (A) Volcano plot of the DAP highlighting the nucleo-adhesome components in red. (B) Venn diagram identifying the A549-meta-nucleo-adhesome components DAP. (C) Dot blot analysis of the co-immunoprecipitation of TGFB1I1 and CSRP2 with HA-KDM6B. (D) Relative quantification of the dots intensity of the dot blot assay were compared with t-tests, result is abbreviated as n.s. for p >0.05 * for p <0.05, ** for p <0.01, *** for p<0.001 and **** for p< 0.0001,, error bars are displayed as SD (E) Immunofluorescence staining of TGFB1I1 and CSRP2 in A549 with or without TGFβ/TNFα treatment. **(F)** PLA between HA-KDM6B and TGFB1I1 or CSRP2 in A549 with or without TGFβ/TNFα treatment. **(G)** Nuclear PLA dots were quantified, and counts were compared between TGFβ/TNFα and untreated conditions with t-tests, result are abbreviated as before

We next focused on two nucleo-adhesome partners of KDM6B, TGFB1I1 and CSRP2, of which Co-IP with KDM6B was confirmed by dot blot (**Figure 3 C and D**). To confirm the ability of CSRP2 and TGFB1I1 to locate to the nucleus, we performed immunofluorescence staining for these proteins in A459 cells under the same treatment (**Figure 3 E)**. Similarly to the observations made by Byron *et al*., TGFB1I1 signal was found in the nucleus, consistent with the western-blot analysis (**Figure 2 B),** even though TGFB1I1 was not part of the identified DAP. Proximity Ligation Assay (PLA) performed between TGFB1I1 and HA-KDM6B or CSRP2 and HA-KDM6B under the same treatment conditions indicated that these interactions were localized to the nucleus and increased during EMT induction (**Figure 3F and G)**. These observations are consistent with the fact that TGFB1I1 and CSRP2 are both induced by TGFβ as previously described(48–51).

### TGFB1I1 and CSRP2 are required for the induction of genes within a KDM6B signature

To investigate the role of KDM6B, TGFB1I1 and CSRP2 during EMT, a knockdown strategy using siRNA against each of these proteins was used. RT-qPCR analysis showed an expected induction of *KDM6B*, mRNA during EMT induction and an siRNA efficiency of 50% extinction (**Figure 4 A**).

RNA-seq analysis after EMT induction identified 7330 Differentially Expressed Genes (DEG) and within them, KDM6B KD clearly shows a signature of genes that depend on KDM6B to be induced during EMT (**Figure 4 B -D**). In fact, 44% of the upregulated DEG after EMT were repressed after KDM6B KD, but only 6% of the repressed genes after EMT were repressed after KDM6B KD.

The next steps were carried out to establish a robust signature of “mesenchymal” genes regulated by KDM6B at the epigenetic level (**Figure 4 E**). First, the 4800 DEG upregulated between treated and untreated conditions with a control siRNA were compared to a database of EMT related genes (dbEMT(52)). This established a set of 386 “mesenchymal” genes in our model. Second, the under-expressed DEG between the KDM6B knockdown versus control siRNA during EMT induction were overlapped with this mesenchymal gene set and with genes showing enriched KDM6B ChIP signal for KDM6B, obtained in the same model from one of our previous publications (10). This overlap showed 1286 genes that exhibited KDM6B ChIP signal as well as under-expression during KDM6B knockdown, 97 of which were part of the previously established model-specific “mesenchymal” gene set.

A second RNAseq analysis was performed in the same conditions but with a knockdown of CSRP2 or TGFB1I1. RT-qPCR analysis showed an induction of *TGFB1I1* and *CSRP2* mRNA during EMT and an siRNA efficiency of around 80%. The ability of the siRNA to lower proteins levels of TGFB1I1 and CSRP2 was also checked by western blot (**Figure 5 A and B**).

**Fig. 5.**
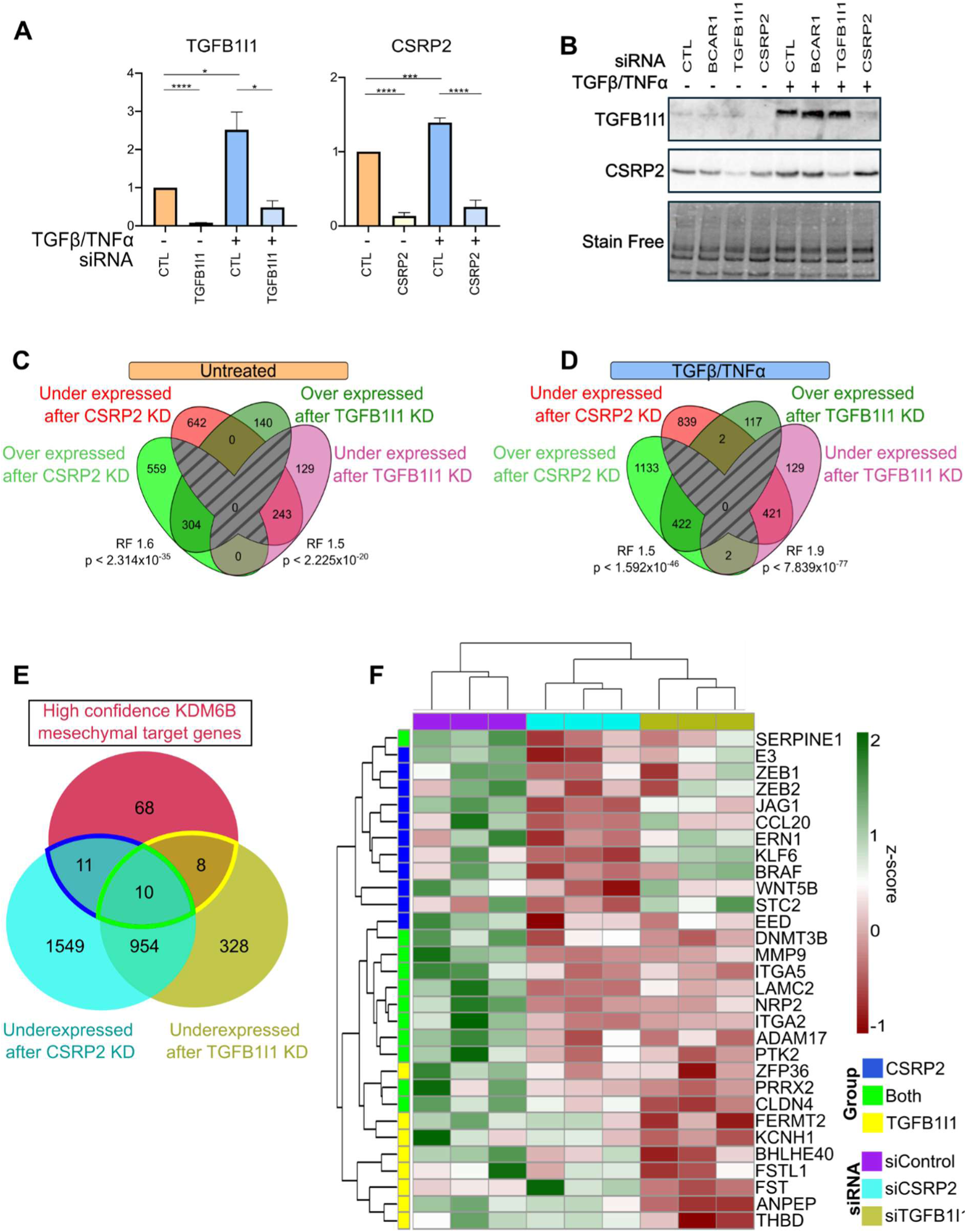
Identifying putative mesenchymal targets of KDM6B-CSRP2 and KDM6B-TGFB1I1 complexes. Knockdown of CSRP2 and TGFB1I1 was performed in A549 cells induced or not in EMT using TGFβ/TNFα then a second RNA-seq analysis was conducted. Efficiency of the siRNA was assessed by (A) RT-qPCR and (B) western blot. Venn diagrams representing how DEG after TGFB1I1 and CSRP2 KD significantly overlap with each other (C) without or (D) with TGFβ/TNFα treatment. (E) Putative mesenchymal targets of KDM6B-TGFB1I1 or KDM6B-CSRP2 complexes were identified by comparing the down regulated genes after TGFB1I1 or CSRP2 KD to the high confidence KDM6B mesenchymal target genes previously identified. (E) The z-scores for these putative targets were then plotted on a heatmap.

Interestingly, TGFB1I1 seemed to share a strong functional relationship with CSRP2 as 243 of the 372 identified downregulated DEG after TGFB1I1 KD were also DEG after CSRP2 KD and 304 of the 444 upregulated DEG after TGFB1I1 KD were also DEG after CSRP2 KD (**Figure 5 C)**. Furthermore, all the identified DEG were upregulated by both KD or down regulated by both KD. A similar observation can be made after EMT induction with most DEG after TGFB1I1 KD also being DEG after CSRP2 KD and all but 4 DEG being upregulated by both KD or down regulated by both KD (**Figure 5 D**).

The repressed DEG identified by this second analysis during EMT induction against the control siRNA were compared to the high confidence KDM6B mesenchymal gene set previously identified (**Figure 5 E and F)**. Amongst the 97 genes of the set, 11 were downregulated during CSRP2 knockdown, 8 during TGFB1I1 knockdown and 10 were downregulated by both knockdowns making them putative targets of a CSRP2-KDM6B or TGFB1I1 complex. In total, almost a third (30%) of KDM6B mesenchymal target genes were co-regulated by CSRP2, TGFB1I1 or both, suggesting that these genes are putative targets of complexes involving KDM6B with one or both partners.

### MMP9, ITGA5 and LAMC2 induction during EMT depends on KDM6B, TGFB1I1 and CSRP2

Based on RNAseq results and literature, three target genes of KDM6B, TGFB1I1 and CSRP2 were selected for further validation. Integrin α-5 (*ITGA5*) and Laminin subunit γ2 (*LAMC2*), two genes downregulated by both CSRP2 and TGFB1I1 knockdown, and Matrix Metalloproteinase-9 (*MMP9*) was also included as it is a previously described mesenchymal target of KDM6B(15) and a CSRP2 regulated gene(47). RT-qPCR results confirmed that *ITGA5*, *LAMC2* and *MMP9* are mesenchymal genes regulated by KDM6B. Their expression was induced by TGFβ/TNFα treatment, and KDM6B knockdown reduced their expression by about 50% during EMT induction (**Figure 6 A-C**). Furthermore, the overexpression of these genes during EMT induction was also found to be dependent on TGFB1I1 and CSRP2 as knockdown of these proteins reduced *ITGA5*, *LAMC2* and *MMP9* expression by about 50% as well. Interestingly, knockdown of KDM6B, TGFB1I1 or CSRP2 also reduced the expression of these genes in the absence of TGFβ/TNFα, suggesting that their role in regulating these genes is not strictly dependent on TGFβ/TNFα signaling. *ITGA5*, *LAMC2* and *MMP9* expression in absence of TGFβ/TNFα induction are indeed non-negligeable as the A549 cell line has been shown to exhibit a hybrid phenotype on the epithelial to mesenchymal phenotype spectrum(53,54).

**Fig. 6.**
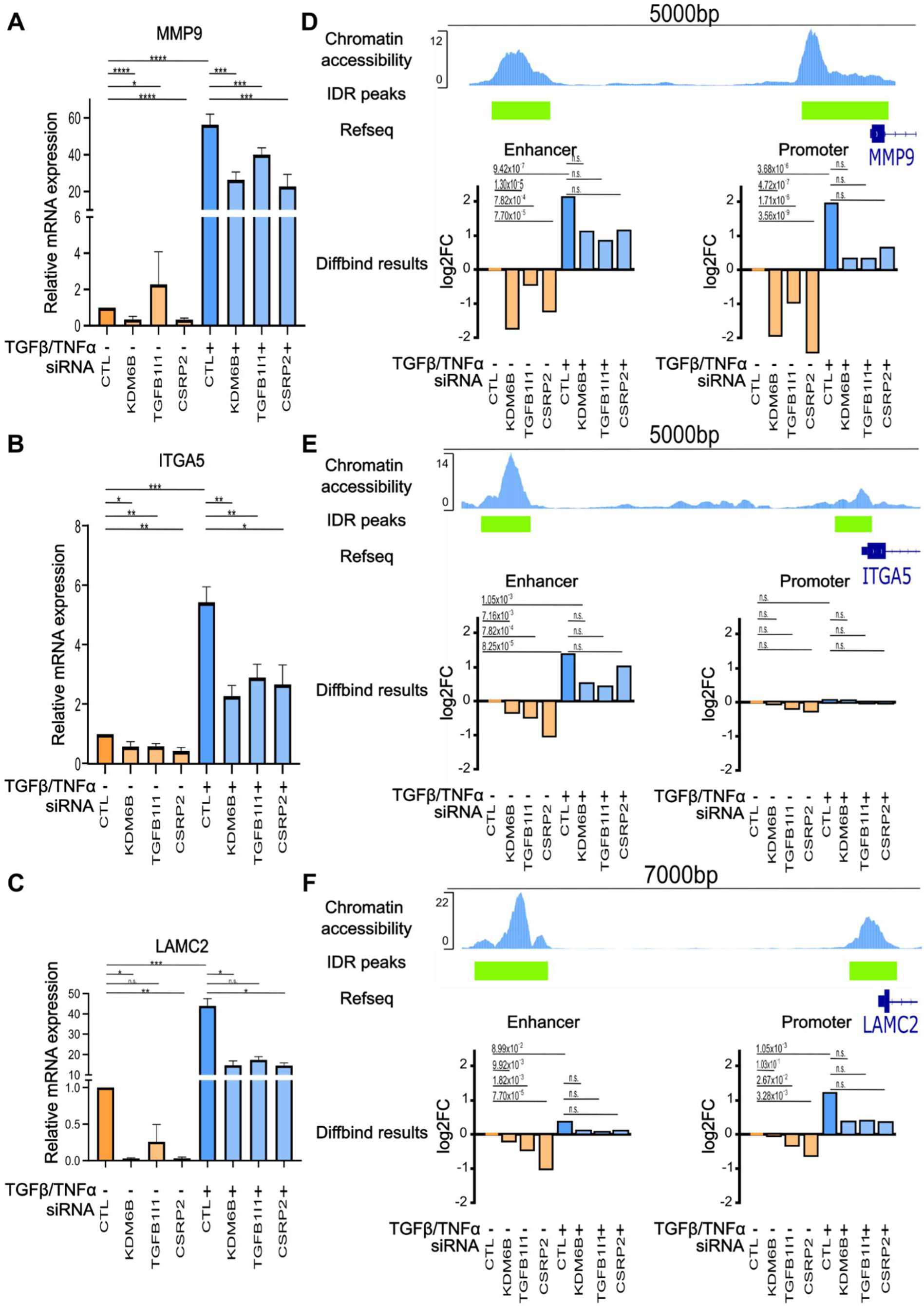
TGFB1I1 and CSRP2 are required for the expression of mesenchymal genes induced at the epigenetic level during EMT. The expression of the three putative target genes *MMP9* (A), *ITGA5* (B) and *LAMC2* (C) mRNA relative to 18S mRNA during EMT induction and knockdown of KDM6B, TGFB1I1 or CSRP2 was confirmed by RT-qPCR. Statistical comparisons were performed with t-tests, results are abbreviated as ns for p >0.05 * for p <0.05, ** for p <0.01, *** for p<0.001 and **** for p< 0.0001, error bars are displayed as SD. The chromatin accessibility of the regulatory regions of *MMP9* (D), *ITGA5* (E) and *LAMC2* (F) following KDM6B, TGFB1I1 or CSRP2 KD or EMT induction was assessed by ATAC-seq. Normalization, fold changes calculation and statistical comparisons were performed using Diffbind, and p-value after Benjamini-Hochberg correction are displayed.

To confirm that these proteins regulate their target genes at the epigenetic level, ATAC-seq was performed (**Figure 6 D-F**). *MMP9* showed decreased accessibility of chromatin in its promoter and enhancer regions after KDM6B and CSRP2 KD, with TGFB1I1 KD having a lesser, but significant effect. In contrast, *LAMC2* showed a significant decrease in chromatin accessibility in its promoter and enhancer after TGFB1I1 and CSRP2 KD but KDM6B KD significantly reduced chromatin accessibility in the enhancer. Lastly, *ITGA5* showed decreased accessibility in its enhancer following all three KD but not in its promoter. All three genes were confirmed to be epigenetically regulated in these regions during EMT induced by TGFβ/TNFα. Interestingly, the ITGA5 promoter did not show increased accessibility after EMT but only in its enhancer. Each of the three genes exhibited a similar pattern of chromatin accessibility after KD and EMT induction which was not found to be significant by statistical analyses.

### Nuclear localization of TGFB1I1 and CSRP2 is associated with a mesenchymal phenotype in lung cancer

To evaluate whether CSRP2 and TGFB1I1 were favorable to establishing a mesenchymal phenotype and migration, wound healing assays were performed (**Figure 7A and B**). CSRP2 and TGFB1I1 knockdowns reduced EMT induced A549 cell migration by about 40%. Next, to assess if CSRP2 and TGFB1I1 are associated with EMT in patients, immunohistochemistry staining of these proteins and VIMENTIN as a mesenchymal marker was performed. As expected, both CSPR2 and TGFB1I1 exhibited an ability to locate to the nucleus or in the cytoplasm (**Figure 7C and D**). Although their respective expression did not increase in EMT positive areas (areas exhibiting high Vimentin expression), CSRP2 and TGFB1I1 nuclear localization was found to be increased in EMT positive tissues (**Figure 7E and F**). In EMT negative areas, 25% of the cells exhibited nuclear staining for TGFB1I1 but in EMT positive areas 50% of the cells exhibited nuclear staining although this difference was not found statistically significant (p =0.1334). For CSRP2 however, a significant difference was observed, with 20% of cells exhibiting nuclear staining in EMT negative areas to 50% in EMT positive areas. These observations are consistent with survival plots obtained with KM plotter(55) as CSRP2 and TGFB1I1 expression did not significantly affect patient prognosis (**Figure 7G and H**). In contrast, high expression of their partner KDM6B as well as their target genes *LAMC2*, *MMP9* and *ITGA5* has previously been associated with significantly lower survival rates(10). Taken together, these results suggest that expression levels of CSRP2 and TGFB1I1 are not responsible for tumor aggressiveness, but their nuclear localization may contribute to it.

**Fig. 7.**
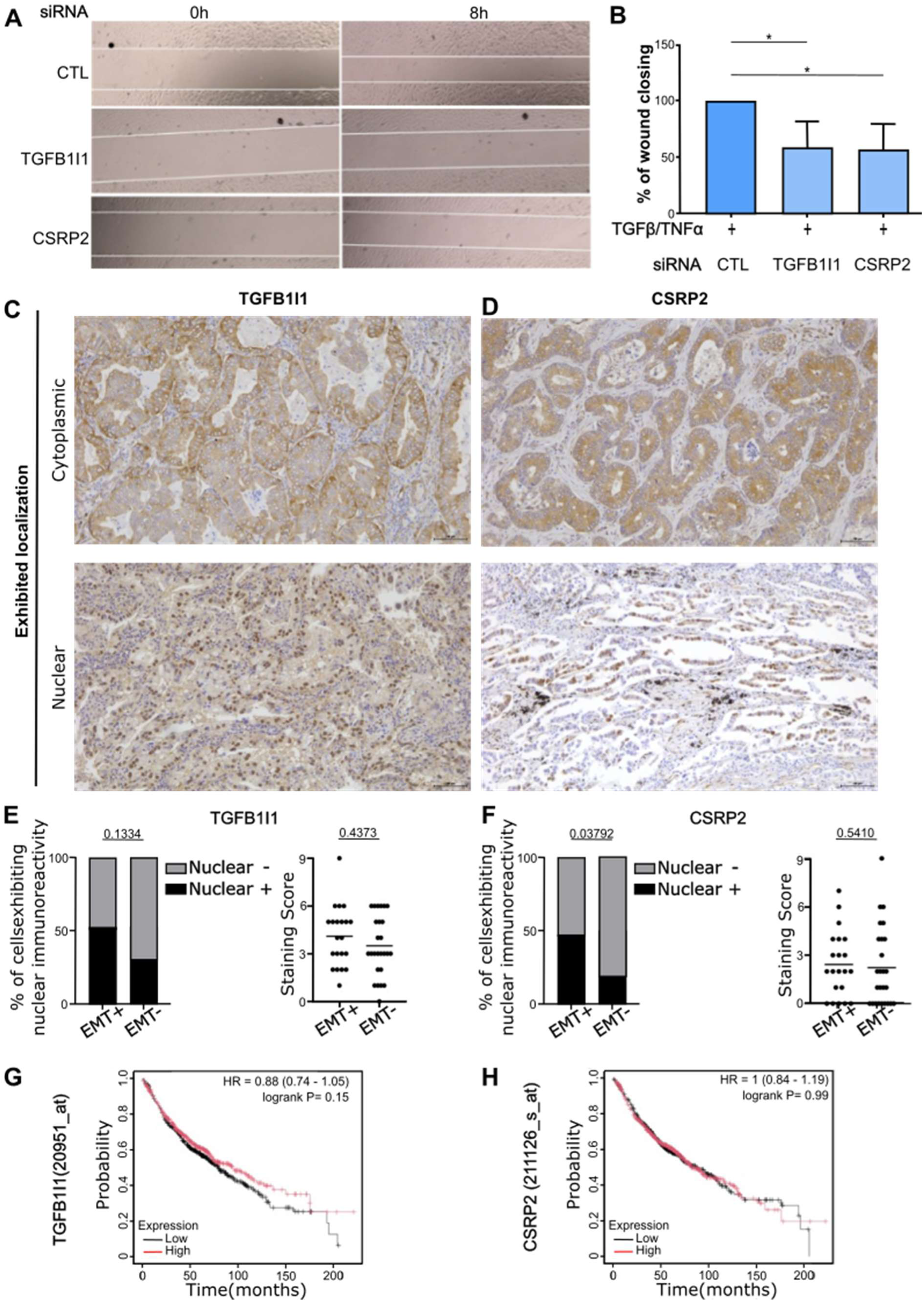
CSRP2 and TGFB1I1 favor a migratory phenotype and shuttle to the nucleus in EMT positive lung cancer tissue. (A) Wound healing assay was performed following TGFβ/TNFα treatment and TGFB1I1 or CSRP2 treatment. Measurements of wound area were performed 0h and 8h after wounding (B) Results are displayed as percent % of wound closing. Statistical comparisons were performed with with t-tests, results are abbreviated as ns for p >0.05 * for p <0.05, ** for p <0.01, *** for p<0.001 and **** for p< 0.0001, error bars are displayed as SD. Examples of IHC staining of TGFB1I1 (C) and CSRP2 (D) showing their ability to localize in the cytoplasm as well as in the nucleus of cancerous cells. Areas of each lung cancer tissues were labelled EMT positive of EMT negative based on Vimentin expression. Quantification of the total staining score as well as percentage of cells exhibiting nuclear immunoreactivity were assessed in EMT negative or EMT positive tissues for TGFB1I1 (E) or CSRP2 (F). Statistical comparisons were performed with t-tests. Kaplan-Meier survival plots for (G) TGFB1I1 and (H) CSRP2 expression in lung cancer obtained with KMplotter.

## DISCUSSION

Both EZH2 and KDM6B have previously been associated with EMT and poor prognosis(1–9). These enzymes epigenetically regulate various genes involved in EMT but the mechanism through which they do so remain unclear, especially for KDM6B. Compared to EZH2, KMD6B has been less studied, and the protein complexes it forms remain poorly characterized, largely due to technical challenges. Our co-IP/MS approach allowed us to establish a list of putative partner proteins for KDM6B involved in the EMT process (**Figure 1**). Identification of most known PRC2 components interacting with EZH2 proved the efficiency of our method. For EZH2 as well as KDM6B, partner proteins involved in cell adhesion were identified. These proteins could serve a dual function, acting both in focal adhesions and in the nucleus to regulate EMT. These protein partners were described as part of a broader network of interacting proteins called nucleo-adhesome. Subcellular fractionation and LC-MS/MS showed that many of the nucleo-adhesome components were differentially represented in the nucleus during EMT induction in A549 cells (**Figure 2**). Within the putative partners of KDM6B belonging to the nucleo-adhesome, CSRP2 and TGFB1I1 were promising targets as they have been reported to exhibit co-regulatory activity and to be associated with EMT. Their nuclear localization and interaction with KDM6B were confirmed by immunofluorescence and PLA, respectively. To better understand the role of these interactions, a signature set of mesenchymal genes epigenetically regulated by KDM6B during EMT was established. This signature associated previously published ChIPseq data and RNAseq data in the same model with or without KDM6B knockdown (**Figure 3**). RNAseq analysis revealed about one third of these targets were regulated by TGFB1I1, CSRP2 or both when knocking down these proteins in the same model. Considering the nuclear localization of the identified interactions, the way the target genes *MMP9*, *LAMC2* and *ITGA5* were established using KDM6B ChIPseq, and the change in chromatin accessibility demonstrated in their regulatory regions after KDM6B, TGFB1I1 or CSRP2 knockdown, it is likely that the regulation of these genes by CSRP2 and TGFB1I1 involves epigenetics and KDM6B. However, these results do not provide formal proof of such regulation. Although technically challenging, ChIP experiments would be required to determine whether these interactions are directly involved in regulating H3K27me3. Nevertheless, CSRP2 and TGFB1I1 appear to be necessary to establish some mesenchymal characteristics during EMT, as their knockdown reduced the expression of mesenchymal genes and diminished cell migration (**Figure 4 and 5**). Moreover, the nuclear localization of these proteins appears to be associated with EMT as exhibited by the IHC results (**Figure 5**). To our knowledge, this work is the first, to involve the nucleo-adhesome in a biological context since it was first described. However, the precise interactome and organization of the nucleo-adhesome during EMT remain to be elucidated with proximity-labeling techniques such as BioID. Further research should focus on designing models capable of better sorting what are the specific nuclear roles of CSRP2, TGFB1I1 and other nucleo-adhesome components. Given that CSRP2 and TGFB1I1 act as co-regulators, investigating the transcription factors they interact with to regulate EMT would be of interest. The transcription factor SRF has been shown to interact with CSRP2 to regulate *MMP9* expression and smooth muscle differentiation(46,47).The *ITGA5* promoter has also been shown to be bound and regulated by SRF(56,57). SRF binding sites were also predicted by JASPAR in the promoter of *LAMC2* which makes SRF an interesting lead to further detail the function of a KDM6B-CSRP2 complex in gene regulation during EMT.

## Acknowledgments

French Proteomic Infrastructure (ProFI) project (grant ANR-10-INBS-08 & ANR-24-INBS-0015).

MGX (Montpellier Genomix). Unité Mixte de Service BioCampus Montpellier (UMS 3426 CNRS-INSERM-UM1-UM2

The DImaCell facility, Univ. Bourgogne Franche-Comté, F-25000 Besançon, France.

## Supplementary figures

**Sup. Fig. 1.**
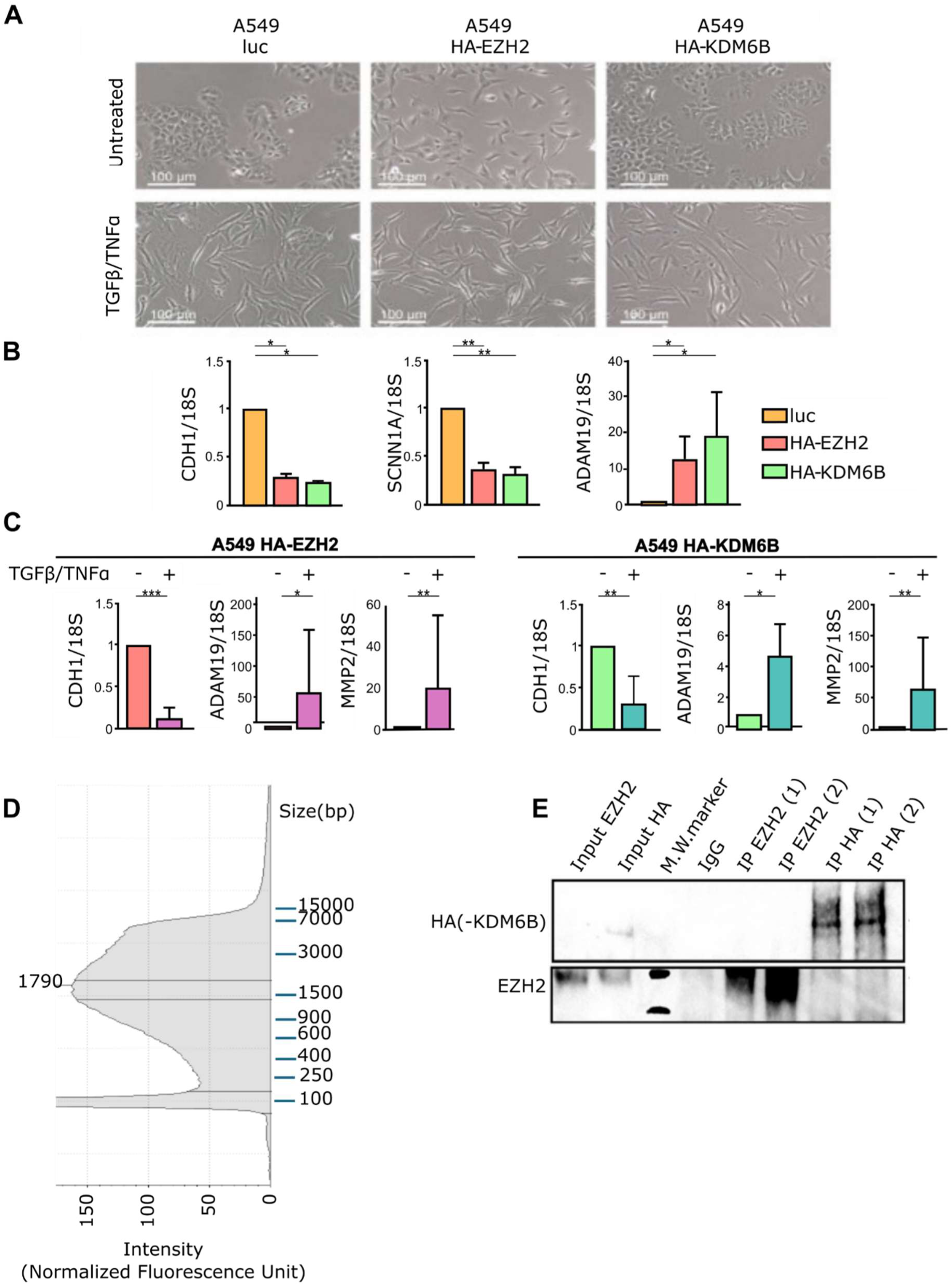
Cell models and co-IP protocol validations. (A) Phase contrast microscopy images of A549 expressing luciferase, HA-EHZ2 or HA-KDM6B with or without TGFβ/TNFα treatment. (B) For each cell line, the expression of epithelial markers CDH1, SCNN1A and mesenchymal marker ADAM19 mRNAs relative to 18S mRNA were assessed by RT-qPCR. (C) In the same way, the expression of epithelial marker CDH1 and mesenchymal markers MMP2 and MMP9 was assessed in A549 HA-EZH2 and A549 HA-KDM6B cells treater or not with TGFβ/TNFα Statistical comparisons were performed with t-tests, results are abbreviated as ns for p >0.05 * for p <0.05, ** for p <0.01, *** for p<0.001 and **** for p< 0.0001, error bars are displayed as SD. (D) Tapestation fragment size profile of chromatin following shearing with our co-immunoprecipitation protocol. (E) Western blot assessing the efficiency of the immunoprecipitation of EZH2 or KDM6B with our co-immunoprecipitation protocol.

**Sup. Fig. 2.**
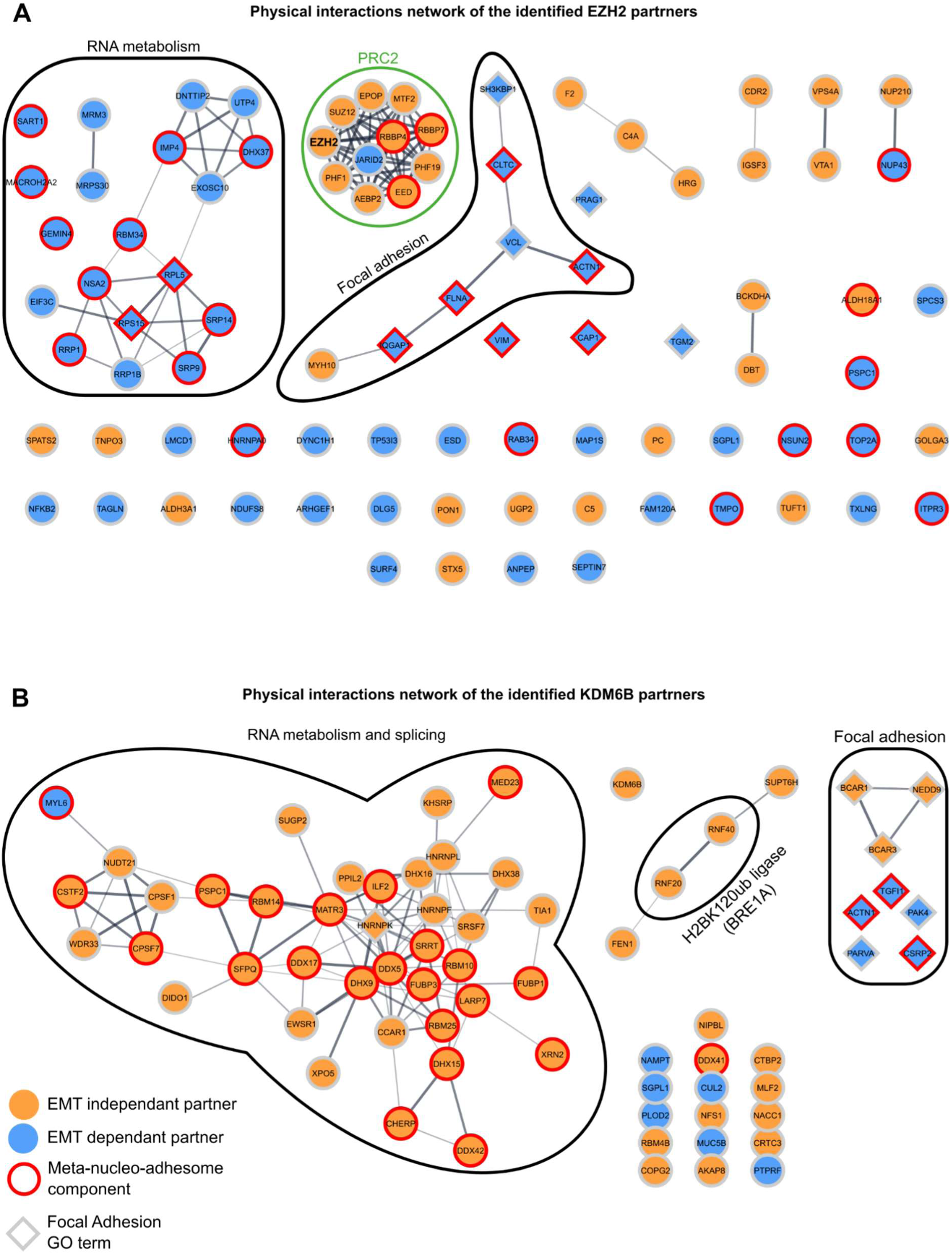
Protein-protein interaction networks for the identified partners of EZH2 and KDM6B. Protein-protein interaction networks were generated by string-db and vizualized using cytoscape for the identified partners of EZH2 (A) and KDM6B (B)

**Sup. Fig. 3.**
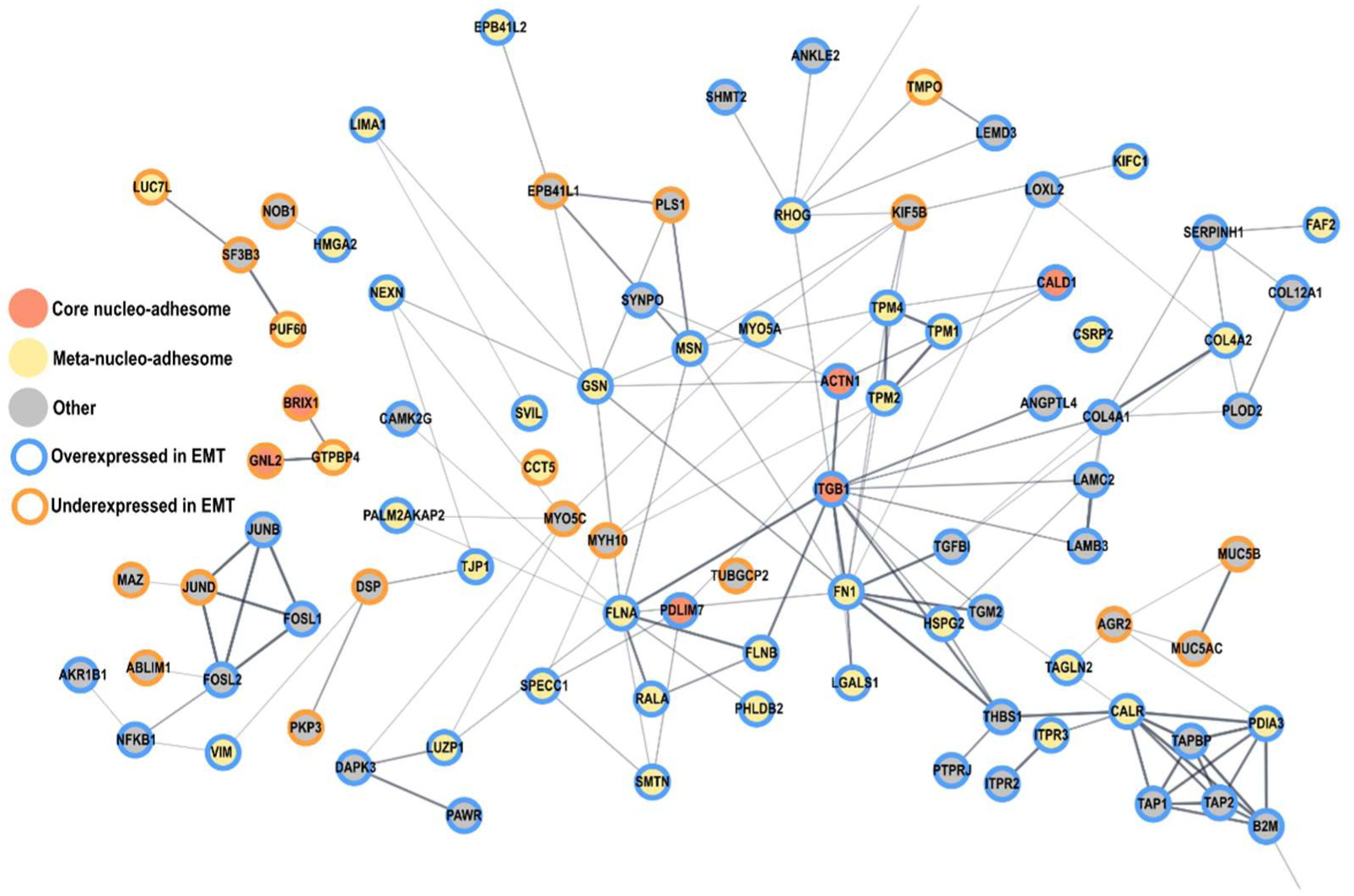
Protein-protein interaction networks of the proteins identified in the nuclear fraction. Protein-protein interaction networks were generated by string-db and visualized using cytoscape for the identified proteins of the nuclear sub-proteome of A549 cells

## REFERENCES

1. Lee HW, Choe M. Expression of EZH2 in renal cell carcinoma as a novel prognostic marker. Pathol Int. nov 2012;62(11):735–41.

2. Xu B, Abourbih S, Sircar K, Kassouf W, Mansure JJ, Aprikian A, et al. Enhancer of zeste homolog 2 expression is associated with metastasis and adverse clinical outcome in clear cell renal cell carcinoma: a comparative study and review of the literature. Arch Pathol Lab Med. oct 2013;137(10):1326–36.

3. Bachmann IM, Halvorsen OJ, Collett K, Stefansson IM, Straume O, Haukaas SA, et al. EZH2 expression is associated with high proliferation rate and aggressive tumor subgroups in cutaneous melanoma and cancers of the endometrium, prostate, and breast. J Clin Oncol Off J Am Soc Clin Oncol. 10 janv 2006;24(2):268–73.

4. Behrens C, Solis LM, Lin H, Yuan P, Tang X, Kadara H, et al. EZH2 protein expression associates with the early pathogenesis, tumor progression, and prognosis of non-small cell lung carcinoma. Clin Cancer Res Off J Am Assoc Cancer Res. 1 déc 2013;19(23):6556–65.

5. Tang B, Qi G, Tang F, Yuan S, Wang Z, Liang X, et al. Aberrant JMJD3 Expression Upregulates Slug to Promote Migration, Invasion, and Stem Cell-Like Behaviors in Hepatocellular Carcinoma. Cancer Res. 15 nov 2016;76(22):6520–32.

6. Li Q, Hou L, Ding G, Li Y, Wang J, Qian B, et al. KDM6B induces epithelial-mesenchymal transition and enhances clear cell renal cell carcinoma metastasis through the activation of SLUG. Int J Clin Exp Pathol. 2015;8(6):6334–44.

7. Cregan S, Breslin M, Roche G, Wennstedt S, MacDonagh L, Albadri C, et al. Kdm6a and Kdm6b: Altered expression in malignant pleural mesothelioma. Int J Oncol. mars 2017;50(3):1044–52.

8. Liang S, Yao Q, Wei D, Liu M, Geng F, Wang Q, et al. KDM6B promotes ovarian cancer cell migration and invasion by induced transforming growth factor-β1 expression. J Cell Biochem. janv 2019;120(1):493–506.

9. Lachat C, Boyer-Guittaut M, Peixoto P, Hervouet E. Epigenetic Regulation of EMT (Epithelial to Mesenchymal Transition) and Tumor Aggressiveness: A View on Paradoxical Roles of KDM6B and EZH2. Epigenomes. 20 déc 2018;3(1):1.

10. Lachat C, Bruyère D, Etcheverry A, Aubry M, Mosser J, Warda W, et al. EZH2 and KDM6B Expressions Are Associated with Specific Epigenetic Signatures during EMT in Non Small Cell Lung Carcinomas. Cancers. 5 déc 2020;12(12):E3649.

11. Sample RA, Nogueira MF, Mitra RD, Puram SV. Epigenetic regulation of hybrid epithelial-mesenchymal cell states in cancer. Oncogene. juill 2023;42(29):2237–48.

12. Lin YT, Wu KJ. Epigenetic regulation of epithelial-mesenchymal transition: focusing on hypoxia and TGF-β signaling. J Biomed Sci. 2 mars 2020;27(1):39.

13. Sun L, Fang J. Epigenetic regulation of epithelial–mesenchymal transition. Cell Mol Life Sci CMLS. 8 juill 2016;73(23):4493–515.

14. Wu MZ, Tsai YP, Yang MH, Huang CH, Chang SY, Chang CC, et al. Interplay between HDAC3 and WDR5 Is Essential for Hypoxia-Induced Epithelial-Mesenchymal Transition. Mol Cell. 2 sept 2011;43(5):811–22.

15. Peixoto P, Etcheverry A, Aubry M, Missey A, Lachat C, Perrard J, et al. EMT is associated with an epigenetic signature of ECM remodeling genes. Cell Death Dis. 27 2019;10(3):205.

16. Lachat C, Boyer-Guittaut M, Peixoto P, Hervouet E. Epigenetic Regulation of EMT (Epithelial to Mesenchymal Transition) and Tumor Aggressiveness: A View on Paradoxical Roles of KDM6B and EZH2. Epigenomes. mars 2019;3(1):1.

17. Sui A, Xu Y, Yang J, Pan B, Wu J, Guo T, et al. The histone H3 Lys 27 demethylase KDM6B promotes migration and invasion of glioma cells partly by regulating the expression of SNAI1. Neurochem Int. 1 mars 2019;124:123–lp.

18. Lee SH, Kim O, Kim HJ, Hwangbo C, Lee JH. Epigenetic regulation of TGF-β-induced EMT by JMJD3/KDM6B histone H3K27 demethylase. Oncogenesis. 26 févr 2021;10(2):17.

19. Oktyabri D, Tange S, Terashima M, Ishimura A, Suzuki T. EED regulates epithelial-mesenchymal transition of cancer cells induced by TGF-β. Biochem Biophys Res Commun. 10 oct 2014;453(1):124–30.

20. Tange S, Oktyabri D, Terashima M, Ishimura A, Suzuki T. JARID2 is involved in transforming growth factor-beta-induced epithelial-mesenchymal transition of lung and colon cancer cell lines. PloS One. 2014;9(12):e115684.

21. Lee SH, Kim O, Kim HJ, Hwangbo C, Lee JH. Epigenetic regulation of TGF-β-induced EMT by JMJD3/KDM6B histone H3K27 demethylase. Oncogenesis. 26 févr 2021;10(2):17.

22. Mohammed H, Taylor C, Brown GD, Papachristou EK, Carroll JS, D’Santos CS. Rapid immunoprecipitation mass spectrometry of endogenous proteins (RIME) for analysis of chromatin complexes. Nat Protoc. févr 2016;11(2):316–26.

23. Byron A, Griffith BGC, Herrero A, Loftus AEP, Koeleman ES, Kogerman L, et al. Characterisation of a nucleo-adhesome. Nat Commun. 1 juin 2022;13:3053.

24. Bouyssié D, Hesse AM, Mouton-Barbosa E, Rompais M, Macron C, Carapito C, et al. Proline: an efficient and user-friendly software suite for large-scale proteomics. Bioinformatics. 15 mai 2020;36(10):3148–55.

25. Wieczorek S, Combes F, Lazar C, Giai Gianetto Q, Gatto L, Dorffer A, et al. DAPAR & ProStaR: software to perform statistical analyses in quantitative discovery proteomics. Bioinformatics. 1 janv 2017;33(1):135–6.

26. Stirling DR, Swain-Bowden MJ, Lucas AM, Carpenter AE, Cimini BA, Goodman A. CellProfiler 4: improvements in speed, utility and usability. BMC Bioinformatics. 10 sept 2021;22(1):433.

27. Bolger AM, Lohse M, Usadel B. Trimmomatic: a flexible trimmer for Illumina sequence data. Bioinforma Oxf Engl. 1 août 2014;30(15):2114–20.

28. Bray NL, Pimentel H, Melsted P, Pachter L. Near-optimal probabilistic RNA-seq quantification. Nat Biotechnol. mai 2016;34(5):525–7.

29. Love MI, Huber W, Anders S. Moderated estimation of fold change and dispersion for RNA-seq data with DESeq2. Genome Biol. 5 déc 2014;15(12):550.

30. Jacquet M, Hervouet E, Baudu T, Herfs M, Parratte C, Feugeas JP, et al. GABARAPL1 Inhibits EMT Signaling through SMAD-Tageted Negative Feedback. Biology. 24 sept 2021;10(10):956.

31. Bruyere D, Monnien F, Colpart P, Roncarati P, Vuitton L, Hendrick E, et al. Treatment algorithm and prognostic factors for patients with stage I–III carcinoma of the anal canal: a 20-year multicenter study. Mod Pathol. 1 janv 2021;34(1):116–30.

32. Hubert P, Herman L, Roncarati P, Maillard C, Renoux V, Demoulin S, et al. Altered α-defensin 5 expression in cervical squamocolumnar junction: implication in the formation of a viral/tumour-permissive microenvironment. J Pathol. 2014;234(4):464–77.

33. Liu Z wei, Zhang Y ming, Zhang L ying, Zhou T, Li Y yang, Zhou G cheng, et al. Duality of Interactions Between TGF-β and TNF-α During Tumor Formation. Front Immunol. 5 janv 2022;12:810286.

34. Kasai H, Allen JT, Mason RM, Kamimura T, Zhang Z. TGF-beta1 induces human alveolar epithelial to mesenchymal cell transition (EMT). Respir Res. 9 juin 2005;6(1):56.

35. Yamauchi Y, Kohyama T, Takizawa H, Kamitani S, Desaki M, Takami K, et al. Tumor necrosis factor-alpha enhances both epithelial-mesenchymal transition and cell contraction induced in A549 human alveolar epithelial cells by transforming growth factor-beta1. Exp Lung Res. févr 2010;36(1):12–24.

36. Asgarova A, Asgarov K, Godet Y, Peixoto P, Nadaradjane A, Boyer-Guittaut M, et al. PD-L1 expression is regulated by both DNA methylation and NF-kB during EMT signaling in non-small cell lung carcinoma. Oncoimmunology. 2018;7(5):e1423170.

37. Agirre E, Oldfield AJ, Bellora N, Segelle A, Luco RF. Splicing-associated chromatin signatures: a combinatorial and position-dependent role for histone marks in splicing definition. Nat Commun. 29 janv 2021;12(1):682.

38. Bhat P, Chow A, Emert B, Ettlin O, Quinodoz SA, Strehle M, et al. Genome organization around nuclear speckles drives mRNA splicing efficiency. Nature. mai 2024;629(8014):1165–73.

39. Segelle A, Núñez-Álvarez Y, Oldfield AJ, Webb KM, Voigt P, Luco RF. Histone marks regulate the epithelial-to-mesenchymal transition via alternative splicing. Cell Rep. févr 2022;38(7):110357.

40. Chodankar R, Wu DY, Schiller BJ, Yamamoto KR, Stallcup MR. Hic-5 is a transcription coregulator that acts before and/or after glucocorticoid receptor genome occupancy in a gene-selective manner. Proc Natl Acad Sci U S A. 18 mars 2014;111(11):4007–12.

41. Heitzer MD, DeFranco DB. Hic-5, an adaptor-like nuclear receptor coactivator. Nucl Recept Signal. 7 juill 2006;4:e019.

42. Shibanuma M, Kim-Kaneyama J ri, Ishino K, Sakamoto N, Hishiki T, Yamaguchi K, et al. Hic-5 Communicates between Focal Adhesions and the Nucleus through Oxidant-Sensitive Nuclear Export Signal. Yamamoto KR, éditeur. Mol Biol Cell. mars 2003;14(3):1158–71.

43. Kim-Kaneyama J ri, Lei XF, Arita S, Miyauchi A, Miyazaki T, Miyazaki A. Hydrogen peroxide-inducible clone 5 (Hic-5) as a potential therapeutic target for vascular and other disorders. J Atheroscler Thromb. 2012;19(7):601–7.

44. Hoffmann C, Mao X, Dieterle M, Moreau F, Al Absi A, Steinmetz A, et al. CRP2, a new invadopodia actin bundling factor critically promotes breast cancer cell invasion and metastasis. Oncotarget. 22 mars 2016;7(12):13688–705.

45. Chen L, Long X, Duan S, Liu X, Chen J, Lan J, et al. CSRP2 suppresses colorectal cancer progression *via* p130Cas/Rac1 axis-meditated ERK, PAK, and HIPPO signaling pathways. Theranostics. 2 sept 2020;10(24):11063–79.

46. Chang DF, Belaguli NS, Iyer D, Roberts WB, Wu SP, Dong XR, et al. Cysteine-rich LIM-only proteins CRP1 and CRP2 are potent smooth muscle differentiation cofactors. Dev Cell. janv 2003;4(1):107–18.

47. Mgrditchian T, Brown-Clay J, Hoffmann C, Müller T, Filali L, Ockfen E, et al. Actin cytoskeleton depolymerization increases matrix metalloproteinase gene expression in breast cancer cells by promoting translocation of cysteine-rich protein 2 to the nucleus. Front Cell Dev Biol. 2023;11:1100938.

48. Wu ML, Chen CH, Lin YT, Jheng YJ, Ho YC, Yang LT, et al. Divergent signaling pathways cooperatively regulate TGFβ induction of cysteine-rich protein 2 in vascular smooth muscle cells. Cell Commun Signal CCS. 28 mars 2014;12:22.

49. Lin DW, Chang IC, Tseng A, Wu ML, Chen CH, Patenaude CA, et al. Transforming Growth Factor β Up-regulates Cysteine-rich Protein 2 in Vascular Smooth Muscle Cells via Activating Transcription Factor 2*. J Biol Chem. 30 mai 2008;283(22):15003–14.

50. Pignatelli J, Tumbarello DA, Schmidt RP, Turner CE. Hic-5 promotes invadopodia formation and invasion during TGF-β–induced epithelial–mesenchymal transition. J Cell Biol. 30 avr 2012;197(3):421–37.

51. Fernandez I, Martin-Garrido A, Zhou DW, Clempus RE, Seidel-Rogol B, Valdivia A, et al. Hic-5 mediates TGFβ-induced adhesion in vascular smooth muscle cells by a Nox4-dependent mechanism. Arterioscler Thromb Vasc Biol. mai 2015;35(5):1198–206.

52. Zhao M, Liu Y, Zheng C, Qu H. dbEMT 2.0: An updated database for epithelial-mesenchymal transition genes with experimentally verified information and precalculated regulation information for cancer metastasis. J Genet Genomics Yi Chuan Xue Bao. 20 déc 2019;46(12):595–7.

53. Kim N, Hwang CY, Kim T, Kim H, Cho KH. A Cell-Fate Reprogramming Strategy Reverses Epithelial-to-Mesenchymal Transition of Lung Cancer Cells While Avoiding Hybrid States. Cancer Res. 15 mars 2023;83(6):956–70.

54. Gemmill RM, Roche J, Potiron VA, Nasarre P, Mitas M, Coldren CD, et al. ZEB1-responsive genes in non-small cell lung cancer. Cancer Lett. 1 janv 2011;300(1):66–78.

55. Győrffy B. Transcriptome-level discovery of survival-associated biomarkers and therapy targets in non-small-cell lung cancer. Br J Pharmacol. févr 2024;181(3):362–74.

56. Leitner L, Shaposhnikov D, Mengel A, Descot A, Julien S, Hoffmann R, et al. MAL/MRTF-A controls migration of non-invasive cells by upregulation of cytoskeleton-associated proteins. J Cell Sci. 15 déc 2011;124(Pt 24):4318–31.

57. Descot A, Hoffmann R, Shaposhnikov D, Reschke M, Ullrich A, Posern G. Negative Regulation of the EGFR-MAPK Cascade by Actin-MAL-Mediated Mig6/Errfi-1 Induction. Mol Cell. 14 août 2009;35(3):291–304.

